# Context-Aware Hierarchical Fusion for Drug Relational Learning

**DOI:** 10.1101/2024.08.06.606750

**Authors:** Yijingxiu Lu, Yinhua Piao, Sangseon Lee, Sun Kim

## Abstract

Drug relational learning, focused on understanding drug-pair relationships within specific contexts of interest, has emerged as a critical area of investigation for its pivotal role in enhancing the efficacy of disease treatment. The nature of drug relationships exhibits significant variations across diverse contexts, such as different types of cancer cell lines. Existing methods often encounter limitations by either neglecting the incorporation of context information or lacking explicit modeling of the intricate connections within drug-drug-context triplets, due to the difficulty in handling heterogeneous relationships between drugs and context. In this study, we present a novel context-aware hierarchical cross-fusion architecture tailored for drug relational learning. By formulating the problem as the label prediction of drug-drug-context triplets, we explicitly calculate all the relations among the triplets. Considering drugs often function as causes and contexts serve as results, our model enhances the learning of intricate drug pair relations hierarchically fusing the information from drug to context through the learned relations. Empirical results across multiple prediction tasks, including synergy, polypharmacy side effects, and drug-drug interactions, highlight the model’s capability to capture essential information relevant to drug relational learning. Notably, our model demonstrates robust predictive performance even in scenarios of heightened contextual complexity, demonstrating its adaptability in learning context-aware drug relations.

**CCS CONCEPTS:** • **Applied computing** → **Bioinformatics**; **Molecular structural biology**; • **Computing methodologies** → **Artificial intelligence**.

**ACM Reference Format:** Yijingxiu Lu, Yinhua Piao, Sangseon Lee, and Sun Kim. 2024. Context-Aware Hierarchical Fusion for Drug Relational Learning. In *Proceedings of Proceedings of the 30th ACM SIGKDD Conference on Knowledge Discovery and Data Mining (BIOKDD ‘24)*. ACM, New York, NY, USA, 11 pages. https://doi.org/10.1145/nnnnnnn.nnnnnnn

## 1 INTRODUCTION

Drug relational learning explores interactions between pairs of drug molecules within specific biological, chemical, or medical contexts, playing a crucial role in various stages of drug development and disease treatment [9]. Co-administration of two or more drugs is a common practice in treating diseases involving complex biological processes due to its high efficacy [5, 12, 24]. However, the chemical and physical reactions between drugs, known as Drug-Drug Interactions (DDI), can alter the intended functionality of drugs [6]. In addition, the combination of drugs may lead to Adverse Drug Reactions (ADR) due to the natural pharmacology of drugs, which is unpredictable during the development stage, contributing significantly to morbidity and mortality [35]. At the clinical stage, determining whether combination therapy exhibits synergistic effects within specific patients to enhance therapeutic efficacy is a challenging aspect faced by precision medicine [2].

However, the path to exploring drug relational learning is fraught with numerous challenges. The significant constraints of time, funding, and manpower make it challenging for researchers to discover all possible interactions between drugs through traditional laboratory-based methods [26]. Moreover, the vast space of drug combinations further renders high-throughput screening and in vitro experiments impractical for exploring the entire drug combination landscape [41]. Additionally, clinical-based exploration of drug combinations not only poses risks of subjecting patients to unnecessary or even harmful treatments but also falls short in adequately probing the drug combination space in the pre-clinical stages [7]. Furthermore, the current clinical-based approaches may lack the depth needed for thorough exploration.

The development of computational methods, such as machine learning models, has opened up possibilities for exploring the complex and extensive space of drug combination relationships. Many computational approaches have been developed and applied to drug relation learning and prediction tasks [29, 30, 32]. Most traditional machine learning methods rely on the assumption of drug similarity, implying that the drugs exhibiting similar chemical structures or biological functions will also exhibit similar side effects [36]. These methods typically take molecular fingerprints or other manually designed features to represent drugs and predict the relationships between drug pairs by calculating their chemical structural similarity [37], which could lead the model bias to a limited representation space while loss on new drug data. In recent years, with the advancement of deep learning, graph-based learning methods have instead increasingly been applied to extract chemical structures automatically from raw molecular graph data, which have shown promising results in DDI prediction tasks [8, 46].

However, two drugs do not necessarily exhibit the same biological activity or pharmacological properties even though they have similar chemical structures. Therefore, methods built on the assumption of drug structural similarity may extract drug features unrelated to the specific prediction task of interest [15, 42]. Besides, the occurrence of drug-drug interactions that are caused by a series of complex biochemical mechanisms, seldom rely solely on individual molecules. Existing drug relation prediction methods often focus solely on extracting information about individual drugs, overlooking the importance of mutual information between drug pairs [39]. Furthermore, accurately extracting the mutual information presented by a pair of drug molecules within specific biological, chemical, or medical semantics also requires the model to grasp the relationship between drugs and contexts [38]. Research on how to enable models to learn useful information for relationship prediction under specific contexts from drug-drug pairs is still lacking.

Learning relationships between drug-drug and drug-context is such crucial for solving tasks of this nature, whereas, existing research has seldom focused on learning these “relations” when predicting whether a drug combination results in specific toxicity, side effects, or exhibits synergy in specific cell lines. In the process of drug relational learning, (1) as drug interactions change across different contexts, how to extract the crucial features from drugs that are conditioned to the given context? (2) as it is a complex biochemical process, is there any simple but robust way to model the problem? (3) drug and context as two heterogeneous entities, how to properly learn the relations between them? These three questions remark on the main challenges in drug relational learning.

To handle the aforementioned challenges, in this work, we propose a context-aware hierarchical cross-fusion model to predict relationships between drugs. **Considering drugs always act as the causes and context as a result in co-administration**, our model constrains the drug relation learned with the given context through a hierarchical architecture. Inspired by the rule of transitivity in the knowledge graph, we take drug-drug-context triplets as input and explicitly learn the relations of drug-drug and drug-context sequentially to ensure the learning of related drug latent representation conditioned with the other drug and given context. Our model demonstrates significantly superior performance compared to baselines across various drug relation databases, both on classification and regression problems. We observe that as the influence of context becomes larger, our model achieves higher performance gains when compared to the best performing baseline model. Besides, we also conduct a series of ablation studies whose results prove the reasonability of our model architecture in drug relational learning.

## 2 RELATED WORKS

### 2.1 Applications of Drug Relational Learning

As mentioned above, drug relational learning plays a crucial role in various aspects of combination therapy research. Instead of monotherapy, combination therapy is widely employed in the treatment of cancer, viral diseases, and fungal infections due to its various advantages, including high efficacy, fewer side effects, and lower required dosages [5, 12, 24]. Identifying synergistic drug pairs for specific cancer types is thus crucial for improving the effectiveness of anticancer treatments. Additionally, predicting adverse events (AE) or adverse drug reactions (ADR) is another significant application of drug relational learning. AE and ADR are unexpected effects occurring at normal drug doses, which could be unpredictable during the drug development process [1]. Previous works heavily rely on quantitative structure-activity relationship (QSAR) for ADR prediction, which may fail in complex scenarios and lack generalization to new drugs [4, 50]. Developing a model that can uncover potential ADRs, even for new drug pairs, is needed. Another application of drug relational learning is drug-drug interaction, which could lead to a chain of undesired reactions, diminishing drug efficacy and even giving rise to hazardous side effects [11]. Identifying possible reactions that a pair of drugs could lead to is also within the scope of drug relational learning in DDI tasks.

### 2.2 State-of-the-Arts in Drug Relational Learning

Existing drug relational learning methods can be naturally categorized into two groups based on whether they utilize context information. Models that incorporate context information mostly leverage gene expression data from cancer cell lines to represent diseases [16, 26, 38]. However, these methods exhibit two main limitations: (1) Although they consider context information, most simply concatenate gene expression data with drug features, lacking a comprehensive understanding of the relationships between the two. (2) While these methods show significant efficacy in predicting drug synergy, their applicability to tasks such as DDI or ADR prediction is limited. On the other hand, the second group of methods that do not use context information, while achieving good performance in predicting a single type of side effect or interaction [23, 46], struggles to handle the large and complex space of context.

Drug relational learning methods can also be categorized into network-based and network-free, based on whether they utilize heterogeneous networks. Network-based methods construct network of drugs and biomedical concepts such as proteins and diseases to predict synergistic effects of co-administered drugs [14, 48] or multi-drug side effects [21, 51]. These methods leverage shared information from biomedical networks, demonstrating strong performance in drug relationship learning tasks. Such methods often require a large amount of interaction data, and they often struggle to extend to relationship prediction between unseen drugs in the network. Network-free approaches leveraging deep learning methods have emerged recently [26, 46]. The majority of them typically employ pre-defined drug features, such as extended connectivity fingerprints (ECFP) [28], or utilize various chemical and physical properties generated through Python packages like ChemoPy [3]. Instead of directly learning task-specific features with models, they rely on professionally crafted features based on prior knowledge. However, this reliance on expert features may not be sensitive to novel drug features, limiting the adaptability and learning capabilities of the model. Additionally, drugs with similar chemical structures do not necessarily exhibit identical biological activities [33]. The utilization of task-specific features learned through models has been demonstrated to enhance the predictive capability of models [23]. Furthermore, several deep learning methods, including attention mechanisms [38] and hypergraphs [20], have been employed in drug relational learning tasks.

## 3 PROBLEM STATEMENT

As mentioned earlier, our focus is on learning context-conditioned latent drug relation representation to achieve robust prediction.

Let *D* = {*d*_1_, *d*_2_, …, *d*_*n*_ } represent a collection of *n* drugs, and *C* = {*c*_1_, *c*_2_, …, *c*_*m*_ } denote a set of*m* contexts. Consider a set of annotated drug-drug-context triplet tuples (*d*_*i*_, *d*_*j*_, *c, y*), where *d*_*i*_, *d* _*j*_ ∈ *D, c* ∈ *C*, and *y* is the target variable belonging to *Y*. Here, *y* is a scalar value, ranging from negative to positive infinity in regression tasks, and taking binary values (0 or 1) in classification tasks.

To develop a drug relational learning model that is versatile across various tasks, we consider the three most popular tasks in disease treatment: **drug-drug synergy, drug-drug polypharmacy side effect**, and **drug-drug interaction** prediction tasks.

- **Drug-Drug Synergy** task predicts whether a pair of drugs *d*_*i*_, *d* _*j*_ exhibit syerngy in a specif cell line *c*.
- **Drug-Drug Polypharmacy Side Effect** task predicts whether a pair of drugs *d*_*i*_, *d* _*j*_ leads to a specific adverse event *c*.
- **Drug-Drug Interaction** task predicts whether a pair of drugs *d*_*i*_, *d* _*j*_ leads to a particular reaction *c*.

In our experiments, we treat each triplet (*d*_*i*_, *d* _*j*_, *c*) as a sample and train the model to predict *P* (*Y* |*d*_*i*_, *d* _*j*_, *c*).

## 4 MOTIVATION AND PRELIMINARY

In the process of drug relational learning:

1. As drug interactions change across different contexts, how to extract the crucial features from drugs that are conditioned to the given context?
2. As it is a complex biochemical process, is there any simple but robust way to model the problem?
3. Drug and context as two heterogeneous entities, how to properly learn the relations between them?

constitute the main challenges of such tasks. In this section, we discuss how we address these challenges by employing the principle of relational learning, leading to the design of a context-aware hierarchical drug relational learning architecture.

### Extracting hidden features conditioned

In relational learning, the relationship *R* between two entities *A* and *B* forms the minimal unit of a knowledge graph [13]. Using the inherent features of two entities to calculate their relationship is the most intuitive approach. However, due to the global and long-range dependencies that could be exhibited between different entities, using inherent features could lead the model bias to pseudo-correlated features when making predictions. The introduction of hidden variables with the assumption that the entities solely depend on these hidden variables in a relation has been proven to be effective in enabling the learning of non-local dependencies [44, 45]. In drug relational learning, we thus apply individual entity encoders for both drugs and context to abstract their non-local task-related hidden representation:

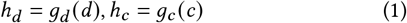

Detailed descriptions are in Section 5.1.

### Learning relationships explicitly

Current drug relational learning usually implicitly learns the relations of drugs and context by feeding the concatenation of their representations into MLPs. One of the main concerns of learning relations like this is the modeling of the relation of drug-drug-context triplet will become vague and less interpretable. To solve this problem, we propose to model the relations of drug-drug and the relations of drug-context independently and explicitly:

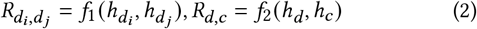

Detailed descriptions are in Section 5.2.

### Handling heterogeneous entities hierarchically

In relational learning, knowledge graphs typically exhibit transitivity [22]. Transitivity also exhibits in drug relational learning. Take DDI for example: if drug *A* alters the function of drug *B*, and the altered function of drug *B* results in toxicity, then the co-administration of drugs *A* and *B* may lead to toxicity. This transitivity indicates that in drug relational learning, drugs could be considered as the causes and context as result, which also implies that the relations of drug-drug and drug-context should be considered sequentially.

Thus, we model the relation of a drug-drug-context triplet by fusing the interaction of drug pairs from drug to drug into the interaction of drug-context pairs from drug to context sequentially in a hierarchical architecture, detailed descriptions are in Section 5.2.

## 5 METHODOLOGY

In this work, inspired by the rule of transitivity in knowledge graph, we take drug-drug-context triplets as input and explicitly learn the relations of drug-drug and drug-context in a hierarchical architecture to ensure the learning of related drug latent representation conditioned with the other drug and given context.

To achieve this, we initially develop uni-entity encoders to independently learn the hidden variables associated with drugs and contexts (**Section 5.1**). These hidden variables are subsequently employed to compute drug-drug and drug-context relations through a hierarchical cross-fusion process (**Section 5.2**). Ultimately, the learned representation is utilized for drug relational learning (**Section 5.3**).

### 5.1 Uni-Entity Encoder

#### 5.1.1 Drug Encoder

Graph-based methods, known for their potent capability to learn topological information, have found extensive application in extracting features from drug data [18, 46]. In this study, we represent drugs using two-dimensional molecular graphs. Specifically, for each drug *d*_*i*_ ∈ *D*, we construct a molecular graph *G*_*i*_ = (*V, E*) based on its SMILES string [40]. Here, *V* = {v_1_, …, v_*N*_ } represents the set of atoms within the molecule, and *E* ⊆ *V* × *V* represents the chemical bonds connecting atoms. We calculate a node feature matrix *X* ∈ ℝ^*N*×*F*^ associated with intermolecular pharmacophores using RDKit [17].

Given a node *v* in the graph of a drug, we employ Graph Isomorphism Network (GIN) [43] to learn node representations *h*_*v*_.

We initialize each node representation using the *x*_*v*_ ∈ ℝ^*F*^ in node feature matrix *X* as:

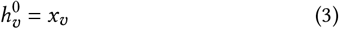

We use *u* ∈ 𝒩 (*v*) to denote the neighboring nodes of the node *v* in a drug 2D graph. At each layer *k*, we update node features according to the following equation:

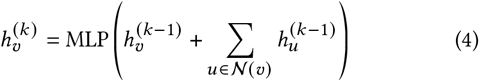

where MLP denotes Multi-Layer Perception, and 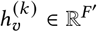 denotes the *k*-th embedding vector associated with node *v*.

We set the total number of layers = 3 and utilize the final node representations for downstream information fusion.

#### 5.1.2 Context Encoder

In different tasks, the term ‘context’ holds different real-world implications. In drug-drug synergy tasks, context represents the cell line for which a drug pair is intended to treat. Models trained specifically for predicting drug-drug synergy often use the gene expression profiles of cancer cell lines as context features. In drug-drug polypharmacy side effects and drug-drug interaction tasks, the context signifies a specific adverse event or reaction type, usually encoded as categorical features. Some models treat the task as a multi-label classification to facilitate learning.

As previously mentioned, in this study, we aim to explore factors influencing drug relational learning and to construct a general model. Therefore, for all three tasks, we utilize categorical feature encoding, specifically, one-hot encoding, as the input for context. Given a context set *C* = {c_1_, …c_*M*_ }, we initialize the feature for each context *c* ∈ *C* and use fully-connected layers to encode the latent context representation as follow:

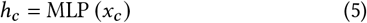

where *x*_*c*_ ∈ ℝ^*M*^ denotes the initialized one-hot encoding feature embedding of context *c*, and *h*_*c*_ ∈ ℝ^*F*′^ represents the hidden embedding of context *c*.

### 5.2 Hierarchical Cross Fusion

In drug relational learning, existing methods often overlook the learning of context information or simply concatenate individually encoded drug and context features before feeding them into a predictor. In this study, we explicitly address the relation between context and drugs to achieve a deeper understanding of the target within drug-context pairs. Recognizing the heterogeneity between drugs and context entities, we introduce a hierarchical structure that sequentially updates homogeneous relations between drugs and heterogeneous relations between drugs and context (Figure 2). Specifically, considering that in reality, drug *A* can alter the function of drug *B*, and drug *B* can similarly alter the function of drug *A*, the interaction between drugs exhibits asymmetry, we decompose the drug-drug relation 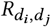 into 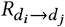 and 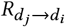. As for drug-context relations 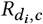 and 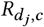, with drugs acting as causes and contexts as effects in biological, chemical, and pharmacological reactions, for practical significance, we only consider the two unidirectional relations from drugs to context, 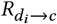 and 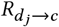.

**Figure 1:**
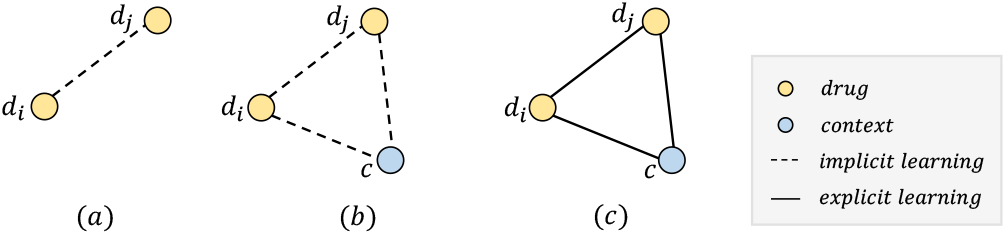
Methods for drug relational learning. Current methods either (a) omit context information or (b) learn the pairwise relations in drug-drug-context triplets implicitly. In this work, we (c) explicitly learn all pair-wise relations in each triplet to improve the robustness of prediction performance.

**Figure 2:**
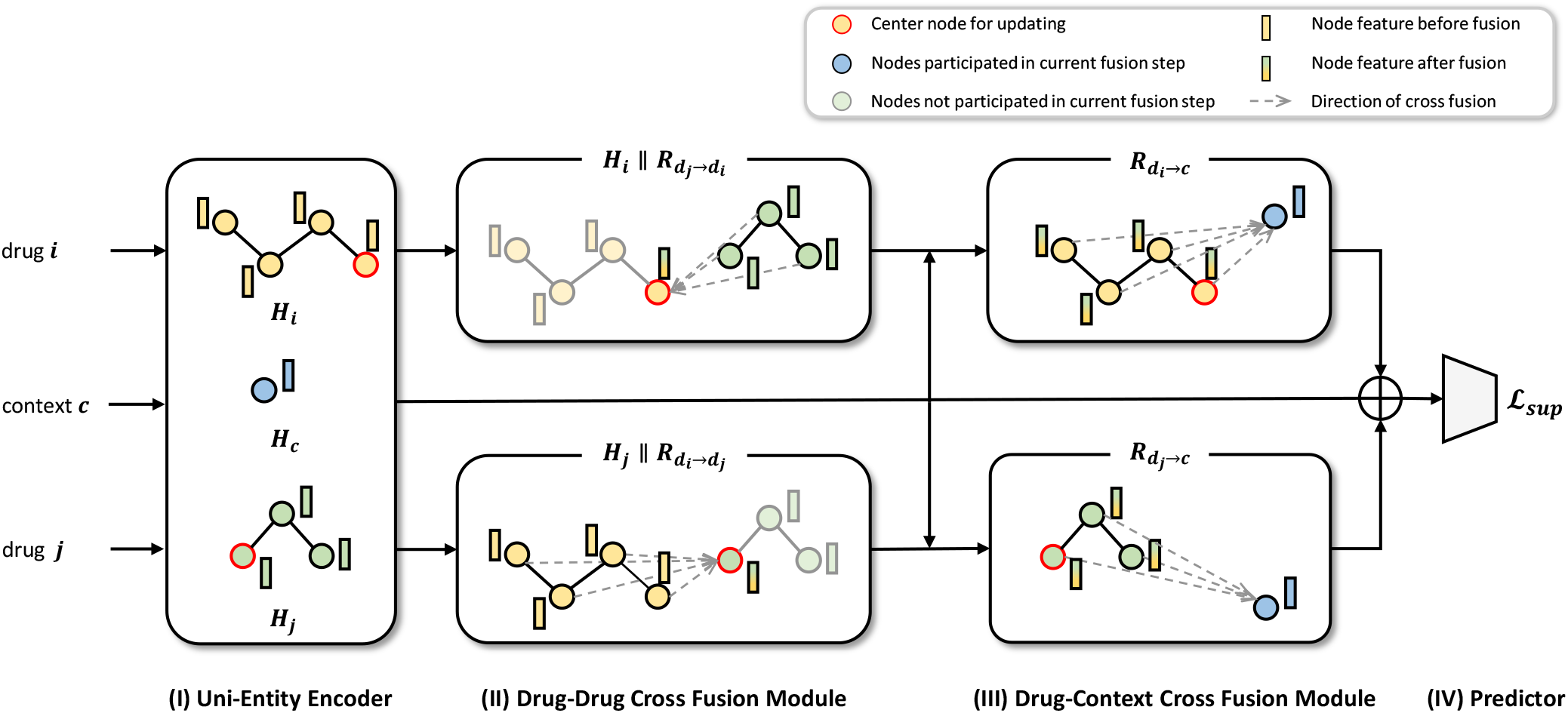
Model architecture. Given a pair of drugs *d*_*i*_ and *d* _*j*_ and the context of interest *c* as input, our architecture predicts whether interaction exists in the certain triplet with hierarchical architecture. *Uni-Entity Encoder* learns the task-related hidden representation of drugs and context independently, after which *Drug-Drug Cross Fusion Module* calculates the possible relations between given drug pairs upon drug hidden representations. Considering drugs act as causes and context (e.g. cancer cell line) act as result in drug relational learning, we constrain the final relation representation with context information by fusing the drug-drug relations with a *Drug-Context Cross Fusion Module*.

Therefore, in drug relational learning, we learn all pairwise relationships within triplets through the following two steps:

- **Drug-Drug Cross Fusion Step** In this step, we compute the relationships 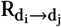 from drug *i* to drug *j* and vice versa. Its practical significance represents any relationship that may exist between a pair of drugs.
- **Drug-Context Cross Fusion Step** In this step, based on the relationships calculated in the previous step between pairs of drugs, we further compute the relationships from drugs to context to constrain the model’s final learning content. Specifically, leveraging the transitivity of relationships, we extend the learned relationship from drug *i* to drug *j* to context in this step. This extension helps extract the portions of the interaction between drug *i* and drug *j* that impact the context, denoted as 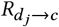. The information from drug *j* to *i* is learned in a similar manner.

#### 5.2.1 Drug-Drug Cross Fusion Module

Inspired by the work of [25], we employ an atom-wise interaction map to calculate the directional relationship 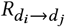 between a pair of drugs *i* and *j*. Specifically, the interaction map is defined as *I*_*ij*_ = sim(*H*_*i*_, *H*_*j*_), where 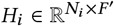 and 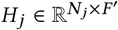 represent the hidden representations of all nodes in drug *i* and drug *j*, and *N*_*i*_ and *N* _*j*_ represent the number of nodes in drug *i* and drug *j*. Consequently, we define 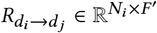 as:

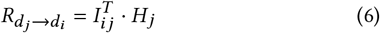

#### 5.2.2 Drug-Context Cross Fusion Module

Based on the relationships computed in Section 5.2.1, we update the representation 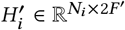 of drug *i* using the following equation:

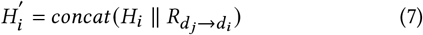

Similarly, we utilize the hidden representations of drugs and the learned hidden representations of context from Section 5.1.2 to compute the relationships between drugs and context:

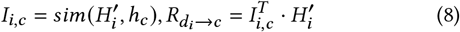

where sim denotes cosine similarity, 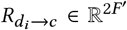 denotes the embedding of drug *i* fused by context *c*. Likewise, 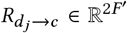 denotes the embedding of drug *j* fused by context *c*.

### 5.3 Triplet Relation Predictor

We obtain the final hidden representation of the drug-drug-context triplet 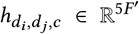, by combining the context hidden representation with the hidden representation of relations that are calculated through the above hierarchical cross-fusion module:

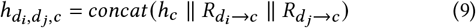

By feeding the hidden representation 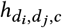 into a single layer of MLP, our model calculates the ultimate output by predicting the probability of a positive label for the drug-drug-context triple.

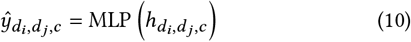

## 6 EXPERIMENTS

### 6.1 Set-up

#### 6.1.1 Dataset

We use six popular benchmark datasets addressing three drug relational learning tasks: combined drug synergy, polypharmacy side effect, and drug-drug interaction. We use four classification tasks from chemicalx [30] and two regression tasks from TDC [11]. Details of datasets are summarized in Tabel 1.

*Four Classification Datasets from Chemicalx:*

- **DrugComb** derived from wet-lab drug combination experiments, containing results from screening studies of drug combinations in various cancer cell lines. It contains 659,333 labels for the synergy between 4,146 drugs across 288 cancer cell lines [47, 49].
- **DrugCombDB** includes experimental data points from highthroughput screening assays of drug combinations, various sources including high-throughput screening and PubMed. It involves 191,391 labels for the synergy between 1,555 drugs across 112 human cell lines [19].
- **TwoSides** incorporates polypharmacy side effects for pairs of drugs, containing 225,070 significant associations between 644 drugs and 10 adverse events, with an equal number of negative samples that were generated without collisions of triples with positive labels [35].
- **DrugBankDDI** contains 86 DDI types, covering 192,284 DDIs from 1,706 drugs. Negative samples were generated without collisions of triples with positive labels [31].

*Two Regression Datasets from TDC:*.

- **DrugComb**^*^ contains 297,098 drug synergy data across 59 NCI-60 cell lines with 129 drugs. It includes four numerical values representing various drug synergies, based on Bliss model, Highest Single Agent (HSA), Loewe additivity model, and Zero Interaction Potency (ZIP) model [27, 47].
- **OncoPolyPharmacology** contains 583 drug-drug combinations tested against 39 human cancer cell lines derived from seven different tissue types. The synergy scores were calculated based on the Loewe additivity values [24, 26].

#### 6.1.2 Baseline models

To comprehensively evaluate our model, we compared it with (1) methods using different drug features and (2) methods with/without learning context information.

##### Models categorized by types of input drug features

In terms of methods that use predefined drug features as input, **MatchMaker** utilized physicochemical features calculated by ChemoPy as drug inputs [3, 16]. **DeepSynergy** additionally employed extended connectivity fingerprints with a radius of 6 (ECFP6) and toxic substructure information [10, 26, 34]. Apart from the above models that use predefined drug features as input, **DeepDrug, SSIDDI, DeepDDS**, and **CMRL** utilized 2D molecular graphs to represent drugs. **DeepDrug** employed GCN, **SSIDDI** used GNN, **DeepDDS** employed GAT and GCN, while **CMRL** used GIN as the drug encoder [18, 23, 38, 46].

##### Models categorized by using or not using context information

Among the above models, only **DeepSynergy, MatchMaker** and **DeepDDS** utilize context information with drug features together as input. Specifically, **DeepSynergy** concatenates drug representations with context representation for prediction directly without using independent encoders. **MatchMaker** generates drug representation conditioned on context information for prediction. **DeepDDS** uses independent encoders to learn hidden representations of drugs and context representations for prediction.

#### 6.1.3 Evaluation protocol

We obtained the reported performances of DeepDrug, DeepSynergy, SSIDDI, MatchMaker, DeepDDS on DrugComb, TwoSides, and DrugBankDDI datasets directly from [30]. For the rest of experiments, we conducted them on an NVIDIA GeForce RTX 3090 to ensure a fair comparison. Each experiment was repeated 5 times with distinct random seeds, ensuring different model parameter initialization and separation of train (80%), validation (10%), and test (10%) sets. Under each setting, our model was trained for 100 epochs with a batch size of 512, learning rate of 1 × 10^−3^, and a dropout rate of 0.2. For fair comparisons, we run the baselines with the same batch size.

#### 6.1.4. Evaluation metrics

In classification tasks, we evaluate the performance of our model using AUROC, AUPRC, and F1. For regression tasks, we assess our model using root mean squared error (RMSE), Spearman’s rank correlation coefficient (SCC), and Pearson’s correlation coefficient (PCC).

#### 6.1.5 Implementation Details

In all the experiments, we apply Adam optimizer for model optimization, and we set the learning rate decreasing on plateau by a factor of 10^−1^ following previous work [18]. For all the tasks, we use a 3-layer GIN [43] as drug encoder, a 2-layer MLP with latent dimension equal to 128 as context encoder, and a single layer of MLP without activation as a final predictor.

### 6.2 Overall Performance

We have compiled the performance of our models on classification tasks in Table 2, and the results for regression tasks are presented in Table 3 and Table 4. Our models consistently outperform the baselines across all tasks. Meanwhile, the majority of baseline models show inconsistent performances across different datasets. For instance, DeepSynergy exhibits better performance on DrugComb and DrugBankDDI, while its performance on DrugCombDB is not good enough. Similarly, CMRL achieves satisfactory results on Drug-CombDB and TwoSides, but its performance on DrugBankDDI is even lower by approximately 2.5% compared to GCN and GAT. These results underscore the effectiveness of our architecture in learning complex drug relations across diverse tasks. In this section, we discuss the advantages of our architecture over baselines by comparing them with several different criteria.

### 6.2.1 Importance of Learning Drug Features in Drug Relational Learning

Among all the baseline models, MatchMaker and DeepSynergy employ drug chemical descriptors calculated by the ChemoPy as inputs, while the remaining methods encode drug features using graph-based techniques. It is observed that the use of predefined drug descriptors can indeed lead to satisfactory model performance on specific datasets. For instance, both MatchMaker and DeepSynergy demonstrated competitive performance with other baselines on DrugBankDDI; however, on other datasets, their performance was less effective compared to graph-based representations.

To address why these methods perform exceptionally well on DrugBankDDI, we conducted a comparison of data statistics across different datasets, as shown in Table 1. It is noteworthy that in the DrugBankDDI dataset, the number of drug-drug pairs is nearly equivalent to the number of drug-drug-context triples. We hypothesize that under such circumstances, where the model can distinguish most drug relations even without knowing context information, well-learned drug representations contribute more to prediction performance. The outstanding performance of our model underscores its ability to learn task-specific latent drug representations. However, models using predefined features are not always the optimal choice, as they may even fall short of vanilla GNN models on certain datasets (DeepSynergy on DrugCombDB, MatchMaker on DrugComb and TwoSides). This emphasizes the significance of learning unbiased hidden features for drug relational learning. The consistently high performance of our model once again highlights the importance of extracting conditioned drug representations.

**Table 1:**
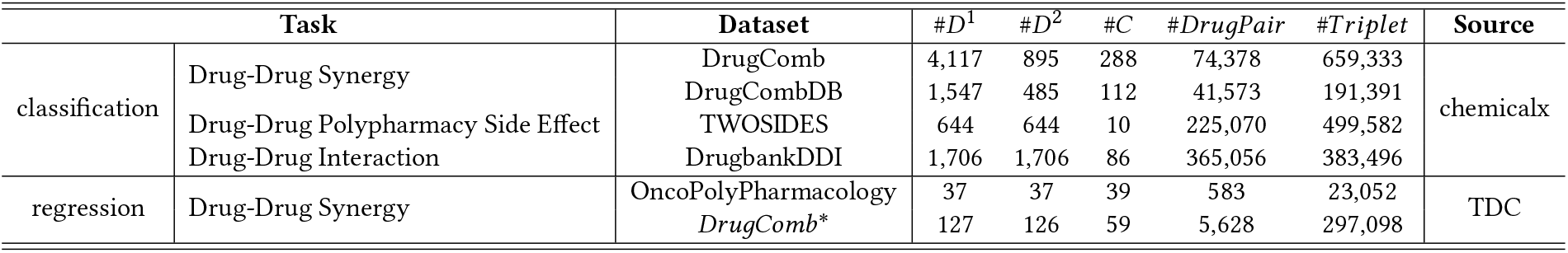
Data Statistics. #*D*^1^: number of drugs in the left. #*D*^2^: number of drugs in the right. #*C*: number of context. #*DrugPair* : number of drug-drug pairs. #*Triplet* : number of drug-drug-context triplets.

**Table 2:** Performance on chemicalx drug relational learning tasks (classification). w/ indicates using context information.

|  | DrugComb |  |  | DrugCombDB |  |  | TwoSides |  |  | DrugBankDDI |  |  |
| --- | --- | --- | --- | --- | --- | --- | --- | --- | --- | --- | --- | --- |
|  | AUROC ↑ | AUPRC ↑ | F1 ↑ | AUROC ↑ | AUPRC ↑ | F1 ↑ | AUROC ↑ | AUPRC ↑ | F1 ↑ | AUROC ↑ | AUPRC ↑ | F1 ↑ |
| GCN (w/) | 66.1 (0.2) | 72.3 (0.2) | 70.5 (0.3) | 76.3 (0.2) | 60.7 (0.4) | 47.3 (0.9) | 92.6 (0.1) | 90.8 (0.2) | 86.1 (0.3) | 97.1 (0.0) | 96.0 (0.0) | 92.5 (0.1) |
| GAT (w/) | 66.2 (0.2) | 72.4 (0.1) | 70.7 (0.6) | 76.1 (0.2) | 60.5 (0.4) | 47.1 (0.6) | 92.4 (0.1) | 90.6 (0.2) | 85.6 (0.5) | 97.0 (0.0) | 95.8 (0.0) | 92.5 (0.0) |
| DeepDrug | 64.3 (0.1) | 70.3 (0.2) | 72.4 (0.1) | 74.0 (0.1) | 57.3 (0.1) | 43.5 (1.0) | 92.3 (0.4) | 90.4 (0.2) | 85.7 (0.2) | 86.1 (0.3) | 82.7 (0.3) | 80.5 (0.2) |
| SSIDDI | 66.3 (0.2) | 72.6 (0.3) | 70.4 (0.3) | 77.9 (0.3) | 62.4 (0.7) | 52.8 (2.8) | 91.7 (0.3) | 89.8 (0.5) | 83.8 (2.9) | 85.5 (0.4) | 82.2 (0.4) | 79.0 (0.5) |
| CMRL | 68.5 (0.2) | 74.8 (0.1) | 70.3 (0.0) | 81.3 (0.1) | 67.0 (0.3) | 58.1 (0.8) | 94.5 (0.0) | 92.9 (0.2) | **91.7 (0.0)** | 94.6 (0.2) | 91.1 (0.0) | 91.3 (0.6) |
| DeepSynergy (w/) | 70.2 (0.3) | 75.8 (0.3) | 72.5 (0.2) | 76.3 (0.5) | 59.8 (0.8) | 48.8 (1.2) | 94.0 (0.1) | 91.9 (0.1) | 88.7 (0.1) | 99.2 (0.1) | 98.7 (0.1) | 96.8 (0.1) |
| MatchMaker (w/) | 66.2 (0.2) | 72.5 (0.1) | 71.2 (0.2) | 78.6 (0.4) | 63.7 (0.9) | 50.8 (2.0) | 91.2 (0.2) | 89.2 (0.1) | 84.9 (0.1) | 98.7 (0.1) | 98.1 (0.1) | 95.9 (0.1) |
| DeepDDS (w/) | 69.1 (0.2) | 74.9 (0.2) | 73.0 (0.3) | 79.0 (0.6) | 64.6 (1.0) | 53.7 (1.3) | 94.1 (0.0) | 92.2 (0.0) | 88.2 (0.2) | 98.9 (0.0) | 98.5 (0.1) | 96.1 (0.1) |
| Ours | **81.8 (0.1)** | **85.7 (0.0)** | **77.8 (0.2)** | **86.0 (0.1)** | **74.8 (0.2)** | **66.4 (0.7)** | **95.6 (0.0)** | **93.4 (0.2)** | 91.6 (0.1) | **99.8 (0.0)** | **99.6 (0.0)** | **99.0 (0.0)** |

**Table 3:** Performance on TDC drug-drug synergy task: drugcomb^*^ (regression). w/ indicates using context information.

|  | ZIP |  |  | HSA |  |  | Bliss |  |  | Loewe |  |  |
| --- | --- | --- | --- | --- | --- | --- | --- | --- | --- | --- | --- | --- |
|  | RMSE ↓ | SCC ↑ | PCC ↑ | RMSE ↓ | SCC ↑ | PCC ↑ | RMSE ↓ | SCC ↑ | PCC ↑ | RMSE ↓ | SCC ↑ | PCC ↑ |
| DeepDrug | 5.35 (0.02) | 0.38 (0.00) | 0.34 (0.01) | 5.88 (0.03) | 0.31 (0.01) | 0.32 (0.00) | 6.02 (0.03) | 0.26 (0.00) | 0.29 (0.00) | 17.18 (0.06) | 0.06 (0.01) | 0.11 (0.01) |
| SSIDDI | 4.56 (0.02) | 0.48 (0.00) | 0.53 (0.00) | 4.93 (0.02) | 0.48 (0.00) | 0.56 (0.01) | 5.13 (0.03) | 0.42 (0.01) | 0.55 (0.00) | 11.52 (0.03) | 0.55 (0.00) | 0.61 (0.00) |
| CMRL | 4.26 (0.01) | 0.49 (0.01) | 0.54 (0.01) | 4.59 (0.01) | 0.50 (0.00) | 0.58 (0.00) | 4.78 (0.01) | 0.45 (0.00) | 0.57 (0.00) | 11.23 (0.03) | 0.57 (0.00) | 0.63 (0.00) |
| DeepSynergy (w/) | 4.02 (0.01) | 0.61 (0.00) | 0.66 (0.00) | 4.38 (0.02) | 0.57 (0.00) | 0.68 (0.00) | 4.55 (0.02) | 0.52 (0.00) | 0.67 (0.00) | 10.63 (0.09) | 0.61 (0.01) | 0.69 (0.00) |
| MatchMaker (w/) | 4.65 (0.02) | 0.51 (0.00) | 0.51 (0.00) | 5.32 (0.03) | 0.42 (0.00) | 0.47 (0.01) | 5.50 (0.03) | 0.38 (0.01) | 0.46 (0.01) | 13.20 (0.31) | 0.39 (0.04) | 0.44 (0.06) |
| DeepDDS (w/) | 4.56 (0.02) | 0.51 (0.00) | 0.54 (0.01) | 5.17 (0.04) | 0.45 (0.00) | 0.53 (0.01) | 5.40 (0.03) | 0.40 (0.01) | 0.51 (0.01) | 12.01 (0.12) | 0.50 (0.01) | 0.58 (0.00) |
| Ours | **3.38 (0.03)** | **0.69 (0.00)** | **0.74 (0.00)** | **3.67 (0.06)** | **0.65 (0.01)** | **0.76 (0.01)** | **3.93 (0.03)** | **0.61 (0.01)** | **0.74 (0.00)** | **9.30 (0.08)** | **0.69 (0.00)** | **0.76 (0.00)** |
| * random split |  |  |  |  |  |  |  |  |  |  |  |  |
| DeepDrug | 5.29 (0.02) | 0.08 (0.01) | 0.12 (0.00) | 5.96 (0.05) | 0.08 (0.01) | 0.10 (0.01) | 6.04 (0.05) | 0.03 (0.02) | 0.06 (0.03) | 14.27 (0.07) | 0.02 (0.01) | 0.02 (0.01) |
| SSIDDI | 6.00 (0.34) | 0.07 (0.04) | 0.07 (0.06) | 7.09 (0.50) | 0.05 (0.02) | 0.05 (0.01) | 7.42 (0.41) | 0.10 (0.05) | 0.07 (0.04) | 19.02 (0.44) | 0.01 (0.01) | 0.01 (0.00) |
| CMRL | 5.36 (0.26) | 0.15 (0.02) | 0.16 (0.02) | 5.29 (0.00) | 0.16 (0.01) | **0.17 (0.01)** | 5.81 (0.38) | 0.12 (0.02) | 0.12 (0.02) | 15.31 (0.44) | 0.03 (0.02) | 0.03 (0.02) |
| DeepSynergy (w/) | 6.02 (0.23) | 0.14 (0.04) | 0.15 (0.05) | 6.72 (0.17) | 0.10 (0.03) | 0.10 (0.05) | 6.76 (0.43) | 0.07 (0.02) | 0.06 (0.03) | 15.28 (0.26) | 0.06 (0.01) | 0.06 (0.01) |
| MatchMaker (w/) | 5.15 (0.02) | **0.22 (0.00)** | 0.21 (0.00) | 5.90 (0.11) | 0.12 (0.02) | 0.12 (0.01) | 5.78 (0.02) | 0.13 (0.01) | 0.10 (0.01) | **13.76 (0.04)** | **0.14 (0.02)** | **0.16 (0.01)** |
| DeepDDS (w/) | 5.12 (0.01) | 0.19 (0.02) | 0.21 (0.03) | **5.70 (0.05)** | 0.09 (0.01) | 0.09 (0.02) | 5.82 (0.01) | 0.05 (0.02) | 0.07 (0.02) | 14.59 (0.29) | 0.05 (0.01) | 0.04 (0.02) |
| Ours | **4.87 (0.11)** | 0.21 (0.00) | **0.23 (0.00)** | 5.27 (0.16) | **0.18 (0.02)** | 0.18 (0.06) | **5.48 (0.12)** | **0.16 (0.04)** | **0.16 (0.05)** | 16.34 (1.70) | 0.05 (0.02) | 0.05 (0.02) |
| * cold drug split |  |  |  |  |  |  |  |  |  |  |  |  |

**Table 4:** Performance on TDC drug-drug synergy task: On-copolyPharmacology (regression). w/ indicates using context information.

|  | OncoPolyPharmacology |  |  |
| --- | --- | --- | --- |
|  | RMSE ↓ | SCC ↑ | PCC ↑ |
| GCN (w/) | 19.86 (1.02) | 0.51 (0.01) | 0.46 (0.02) |
| GAT (w/) | 19.88 (1.04) | 0.51 (0.01) | 0.46 (0.02) |
| DeepDrug | 22.82 (0.69) | 0.44 (0.01) | 0.34 (0.02) |
| SSIDDI | 19.74 (0.76) | 0.54 (0.01) | 0.49 (0.02) |
| CMRL | 19.91 (0.99) | 0.51 (0.01) | 0.45 (0.02) |
| DeepSynergy (w/) | 17.59 (0.59) | 0.65 (0.02) | 0.63 (0.02) |
| MatchMaker (w/) | 18.79 (0.76) | 0.59 (0.01) | 0.56 (0.02) |
| DeepDDS (w/) | 19.75 (0.75) | 0.51 (0.01) | 0.49 (0.01) |
| Ours | **15.28 (0.17)** | **0.73 (0.01)** | **0.74 (0.01)** |

#### 6.2.2 Significance of Using Context Information

In this section, we discuss the significance of incorporating context information. Notably, among all the baseline models, DeepDrug, SSIDDI, and CMRL abstain from utilizing context information, while DeepSynergy, MatchMaker, and DeepDDS incorporate context information into their predictions. We observe that on datasets where the number of drug-drug-context triplets is comparable to the number of drug-drug pairs, such as in TwoSides and DrugBankDDI, models can achieve comparable performance even without considering context information. However, these models exhibit significant performance instability on datasets where drug relations are highly sensitive to different contexts. For instance, CMRL’s performance on DrugBankDDI is approximately 2.5% lower than that of GCN and GAT. In contrast, our model demonstrates robust performance, even as datasets become more complex due to increased contextual variations, underscoring the importance of hierarchical learning of context representation in such tasks.

Particularly noteworthy are the datasets OncoPolyPharmacology and DrugComb^*^, where the ratio of #*Triplets* : #*DrugPair* reaches 39 and 52, respectively. In these tasks, our model consistently outperforms the baseline models across various metrics, demonstrating its promising ability to extract context-aware drug relations.

#### 6.2.3 Contribution of Explicitly Learning Drug Relations

We observed an interesting phenomenon: as the ratio of drug-drug-context triplets to drug-drug pairs increases across different datasets, the performance gain of our model compared to other baselines becomes more significant. As shown in Table 6, with the #*T riplet* : #*DrugPair* ratio increasing from 1.05 to 8.86, the performance gain of our model compared to the highest-performing baseline model (based on AUROC ranking) increased from 0.6% to 11.6%. These results not only demonstrate the successful learning of drug relations by our model, particularly in learning drug relations under specific contexts but also affirm that our model can effectively capture complex drug relations in datasets with significant context influence, maintaining superior performance over other models.

One of the most noteworthy distinctions between our model and other baselines is that our model explicitly learns drug relations through the drug-drug-context triplet, utilizing a hierarchical architecture. To quantitatively analyze the extent to which explicit learning contributes to the overall performance, we modified our architecture by directly concatenating the hidden drug representations and context representation for prediction, ensuring that the relation is implicitly learned through the final MLP layer. The results are presented in the first line of Table 5. It is evident that the model’s performance experiences a significant drop when relations are not explicitly modeled, underscoring the necessity of explicit learning in drug relational learning.

**Table 5:**
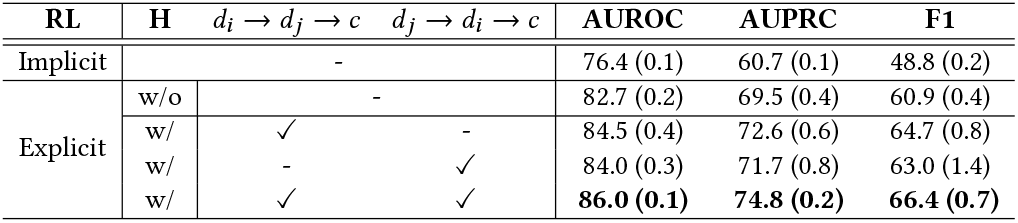
Ablation Study Results on DrugCombDB database. RL: relational learning; H: Hierarchy; *d*_*i*_ → *d* _*j*_ → *c* and *d* _*j*_ → *d*_*i*_ → *c* stand for the direction of relation fusion (Section 5.2).

**Table 6:**
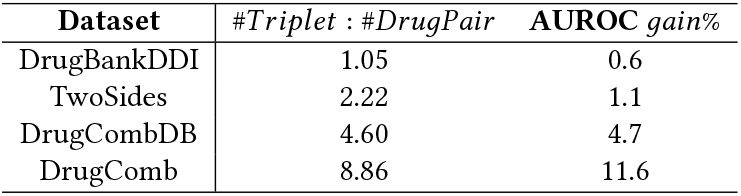
Percent gain of performance of our model when compared with best performed baseline model (AUROC)

#### 6.2.4 Significance of Hierarchical Cross Fusion

Another significant aspect of our model, distinguishing it from the baselines, is the hierarchical cross-fusion architecture. We posit that hierarchically fusing information from drug to drug and subsequently from drug to context assists the model in learning context-constrained features of drug pairs. In tasks like drug relational learning, where the co-administration of drugs is considered, drugs often function as causes, and the contexts, such as cancer cell lines, serve as results.

To assess the extent to which this hierarchical architecture contributes to performance improvement, we modified the architecture into a flat version where both drug-drug relations and drug-context relations are directly learned from the hidden drug representations and context representation. The results in Table 5 indicate that without hierarchy, the model’s performance drops by around 3.3% in AUROC. This suggests that the hierarchical architecture effectively filters out features that are irrelevant to model prediction.

Furthermore, given that our architecture operates under the assumption that the model can learn significant latent representations of each drug related to another under a given context, we also investigate whether the fusion of relations from only one single direction, such as from *d*_*i*_ → *d* _*j*_ → *c* or from *d* _*j*_ → *d*_*i*_ → *c*, is sufficient for drug relational learning. To explore this, we conduct an ablation study using single-sided fusion on the DrugCombDB dataset (Table 5). The results indicate that removing either side of the fusion results in a drop in performance, suggesting that both drugs play an important and irreplaceable role in relational learning.

#### 6.2.5 Cold-Drug Settings

To assess the generalization ability of our model in predicting relationships between unknown drug pairs, we adopted a cold-drug setting by partitioning a small subset of drugs from the original dataset. These drugs, along with their corresponding relationships, formed the test set. The remaining drugs, along with their context-conditioned relationships, constituted the training and validation sets. We then compared the performance of our model against existing baselines in this cold-drug setting.

Tables 3 and 7 present the experimental results under the cold-drug setting. Notably, our model outperformed other models by a significant margin on DrugBankDDI, while achieving performance comparable to the best baseline on DrugComb^*^. We observed that on DrugBankDDI, where a diverse set of drugs is present, most models employing graph-based encoders, including ours, CMRL, and DeepDDS, performed well. This could be attributed to the abundance of different drugs in DrugBankDDI, allowing graphbased models to better learn structural features relevant to relationships. Conversely, DrugComb^*^ contains 5,628 distinct drug pairs but has 186,810 drug-drug-triplet combinations, indicating high context diversity in this database. In such a context-rich environment, the ability of models to learn contextual information is more critical for performance. We observed that, apart from our model, MatchMaker and DeepDDS, which also learn context information, achieved promising results.

**Table 7:** Performance under cold-drug setting on Drug-BankDDI dataset (classification). w/ indicates using context information.

|  | DrugBankDDI |  |  |
| --- | --- | --- | --- |
|  | AUROC ↑ | AUPRC ↑ | F1 ↑ |
| DeepDrug | 67.2 (1.0) | 66.6 (1.3) | 55.4 (1.0) |
| SSIDDI | 66.2 (0.8) | 65.9 (1.3) | 53.0 (4.5) |
| CMRL | 94.0 (0.2) | 90.3 (0.3) | 90.0 (0.2) |
| DeepSynergy (w/) | 90.8 (0.6) | 90.6 (0.5) | 73.1 (2.1) |
| MatchMaker (w/) | 92.8 (0.1) | 93.3 (0.6) | 74.7 (2.5) |
| DeepDDS (w/) | 93.4 (0.5) | 94.1 (0.4) | 85.3 (0.6) |
| HypergraphSynergy (w/) | 91.8 (0.4) | 92.6 (0.5) | 85.7 (0.8) |
| Ours | **99.6 (0.0)** | **99.4 (0.1)** | **98.8 (0.1)** |

### 6.3 Case Study

To investigate the effectiveness of our context-aware hierarchical architecture in explicitly learning context-conditioned drug relations, we visualized the t-SNE embeddings of the last latent layer before the output for the test sets of each dataset. As shown in Figure 3, the context information can be discerned more clearly in the latent layers of our model, indicating that our model adequately learns and considers the context-conditioned drug-drug-relations during prediction, which also suggests the significance of model’s ability to learn and distinguish different contexts in drug relational learning tasks.

**Figure 3:**
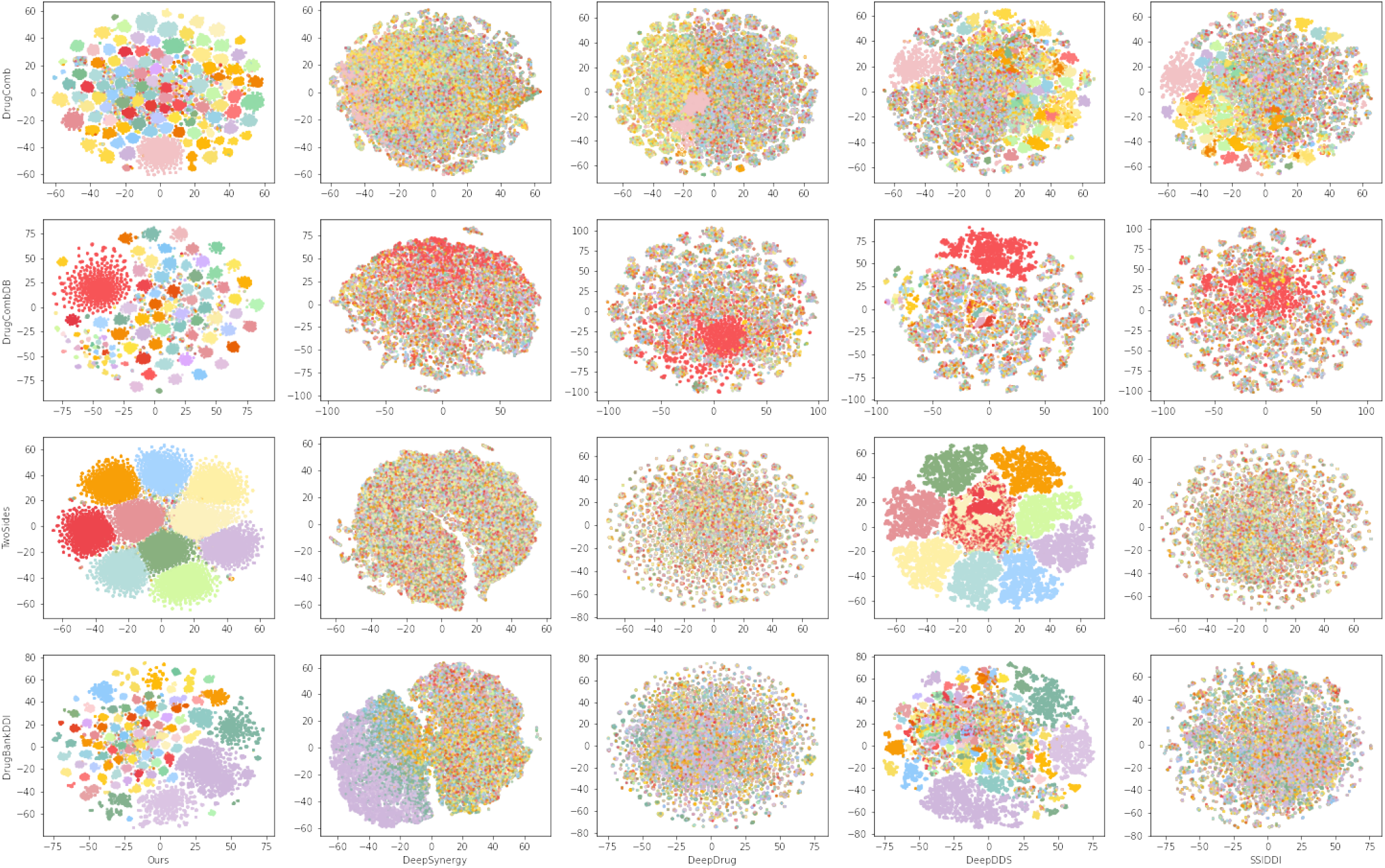
Visualization of model embeddings across different datasets. Each point represents a drug-drug-context triplet, and the color of the scatter corresponds to its unique context label.

## 7 CONCLUSION

Drug relational learning serves as a fundamental approach for comprehending interactions among drug molecules within specific contexts, providing essential insights into drug development and disease treatment. Throughout the drug relational learning process, key challenges include (1) extracting crucial features from drugs conditioned on a given context, (2) modeling the problem in a simple yet robust manner, and (3) appropriately learning the heterogeneous relation between drugs and context. In this study, we tackle the aforementioned challenges in drug relational learning through our proposed context-aware hierarchical cross-fusion model. Recognizing that drugs act as causes and context as a result in co-administration, our model constrains the drug relation learned with context by integrating drug-drug relations into drug-context relations, ensuring the acquisition of pertinent drug latent representations conditioned on both the other drug and context. Our model exhibits significantly superior performance compared to baselines across various drug relation databases, encompassing both classification and regression tasks. Notably, as the influence of context grows, our model achieves even greater performance gains compared to the best-performing baseline model. Analysis of the embedding space further emphasizes the model’s ability to learn context-aware drug relations.

## 8 ACKNWLEDGEMENTS

This research was supported by the Bio & Medical Technology Development Program of the National Research Foundation (NRF) funded by the Ministry of Science & ICT(NRF-2022M3E5F3085677, 2022M3E5F3085681, RS-2023-00257479), Institute of Information & communications Technology Planning & Evaluation (IITP) grant funded by the Korea government(MSIT) [RS-2021–II211343, Artificial Intelligence Graduate School Program (Seoul National University)], funded by AIGENDRUG CO., LTD.. The ICT at Seoul National University provides research facilities for this study.

